# An in-silico approach to identify bioactive phytochemicals from *Houttuynia cordata* Thunb. As potential inhibitors of Human Glutathione Reductase

**DOI:** 10.1101/2023.08.15.553374

**Authors:** Satyam Sangeet, Arshad Khan

## Abstract

Cellular infections underpin the pathogenesis of cancer and malaria. Mitigating cellular oxidative stress via glutathione reductase (GR) inhibition emerges as a promising therapeutic avenue. Exploiting the antioxidant-rich *Houttuynia cordata* Thunb., we investigated natural GR inhibitors. Among 13 docked phytochemicals, Quercetin, Quercitrin, and Sesamin exhibited exceptional GR binding affinities. Molecular Docking analysis highlighted their propensity to precisely target the GR active site. Subsequent 150 ns molecular dynamics simulations corroborated their robust interactions, unveiling dynamic stabilizing effects on the protein structure and bolstering their antioxidant potential. Furthermore, ADME-Tox profiling affirmed their favourable drug-like attributes. These findings underscore *H. cordata’s* reservoir of potent antioxidants, poised to combat various maladies, including malaria and cancer. This study distinctly accentuates the distinctive outcomes and paramount significance of harnessing *H. cordata* phytochemicals as efficacious antioxidants, unravelling novel therapeutic avenues.

## Introduction

Glutathione, a biological antioxidant found in nearly all mammalian tissues, plays a crucial role in maintaining cellular homeostasis. It is a tripeptide made up of; glutamic acid, cysteine, and glycine (Teskey et al. 2018). It exists in the thiol reduced (GSH) and disulphide oxidized form (GSSG). GSH is the most abundant intracellular non-protein thiol, present in millimolar concentrations. GSH accounts for more than 98 percent of the total glutathione and is stored in three main reservoirs in eukaryotic cells. The bulk of cellular GSH (80-85%) is found in the cytosol; 10-15 percent is found in the nucleus and a smaller percentage found in the endoplasmic reticulum and mitochondria. The active thiol group of GSH is found in the cysteine residue and plays a role in antioxidant function either directly by detoxifying reactive oxygen and nitrogen species (ROS and RNS) or indirectly through GSH-dependent peroxidase-catalysed reactions (Couto et al. 2016). As a result, GSH is involved in a number of key detoxification reactions and has a high capacity for preventing oxidative stress-induced cellular damage. Hence, the intracellular redox state, which is characterized by the levels and ratios of oxidized (GSSG) and reduced (GSH) glutathione, is regarded as a significant indication of the cellular health (Lu 2013).

GSH is generated by glutamyl cysteine synthase (GCL) and glutathione synthase (GSS) in a two-step ATP-dependent process. The first step involves GCL forming a peptide bond between glutamate and cysteine, followed by the addition of glycine with the aid of GSS (Couto et al. 2016). While a rise in GSH appears to be a universal cellular response to oxidative stress, decreased GSH levels tend to develop some diseases like mitochondrial diseases (Enns et al. 2014). Conversion of oxidized glutathione (GSSG) to its reduced form (GSSH) is performed by an enzyme Glutathione Reductase (GR) in the presence of NADPH and an FAD prosthetic group. GR is the only enzyme which is a flavoprotein that catalyses the reduction of oxidized glutathione to reduced glutathione. This reaction has the purpose of maintaining a high GSH/GSSG ratio in the cell. In several diseased conditions, the GR levels were found to be higher in patients with a number of tumours, including lung cancer (Rédei 2008). In normal physiological environments, GSH’s primary function is to prevent oxidative damage of the cells by inactivating xenobiotics and oxidants. However, some cells use the glutathione mechanism effectively to function in diseased states; increased glutathione levels and dependent enzymes can reduce therapeutic effectiveness by cleaning the radical molecules. As a result, inhibiting the GR enzyme is an effective therapeutic target in the fight against diseases like malaria and cancer, as it alleviates the levels of GSH (Güller et al. 2021).

Glutathione reductase inhibitors (GRIs) have recently gained popularity in malarial and cancer medications. Steroidal compounds, because of their powerful immunomodulating and anti-inflammatory effects, are now used in a variety of clinical settings. Some corticosteroids - dexamethasone, prednisolone, methylprednisolone and 5-methyl-2,4-dihydro-3H-1,2,4-triazol-3-one’s aryl Schiff base derivatives, N-Methylpyrrole derivatives have been known to inhibit GR (Şentürk and Şentürk 2020; Balaydın et al. 2018; Kocaoğlu et al. 2019). These synthetic drugs however pose major side effects that may prove grave.

There have been several studies conducted to inhibit the GR. For example, the *in-silico* and *in-vitro* studies of the inhibition of GR using phenolic compounds such as curcumin, quercetin and resveratrol was carried out by (Guller et al. 2021) and it has been seen that curcumin is an efficient inhibition of GR in comparison to quercetin and resveratrol. An interesting information to notice is that the phenolic compound quercetin is also present in a plant *Houttuynia cordata* Thunb., along with several other antioxidant phytochemicals (Kumar et al. 2014). This makes *H. cordata* an interesting target to explore as it possesses antioxidant activities (Ju et al. 2021; Nuengchamnong et al. 2009; Tian et al. 2012; Song et al. 2021).

*Houttuynia cordata* Thunb. (HCT) is a flowering and perennial herb native to China, Japan, Korea, and Southeast Asia. It is the only species in the genus *Houttuynia*, which belongs to the *Saururaceae* family. HCT has long been used as a medicinal plant (Chen et al. 2014). HCT has been shown to have anti-cancer activity (Yanarojana et al. 2017). It has been demonstrated that its leaves along with lactic acid bacteria ferments green tea and exerts anti-adipogenic and anti-obesity effects (Wang et al. 2018). HCT also possesses antiviral, anti-oxidant, anti-bacterial, and anti-inflammatory properties (Yanarojana et al. 2017). Chen and group (Chen et al. 2014) demonstrated that both aqueous and methanolic extracts of *Houttuynia cordata* had antioxidant properties in Sprague Dawley rats in which the oxidative stress was produced by feeding frying oil. Ng and group (Ng et al. 2014) investigated the antioxidant properties of *H. cordata* and its ability to protect rats from bleomycin-induced pulmonary fibrosis. In both the normal cells and tumour cells, chemotherapeutics, e.g., various xenobiotics are converted to electrophilic molecules which are combined with GSH and removed out of the body (Guller et al. 2021). As a result, chemotherapeutics become ineffective in killing both the normal cells and tumour cells. But tumour cells need to be killed as they are no longer needed inside the body.

Our strategy centres on targeting the active site of GR using the leading phytochemicals from H. cordata. This aims to hinder the binding of the prosthetic group FAD, effectively interrupting the conversion of GSSG to GSH. Therefore, the current study investigates the use of the phytochemicals from *Houttuynia cordata* Thunb. against glutathione reductase (GR) through *in-silico* methodology to prevent it from taking part in reducing GSSG back to GSH and thereby prohibiting tumour cell survival by creating an extremely stressful environment inside the cells.

## Methodology

### Active Site Identification

The active site inside the co-crystallized complex of glutathione reductase (GR) was predicted in using the CASTp (Computed Atlas of Surface Topography of Proteins) online server (Tian et al., 2018). The aforementioned server received the structural coordinates of GR, and further analysis identified the best parameters, identifying the areas showing an active site pocket’s feature.

### Ligand and Receptor Preparation

The structure of the phytochemicals was downloaded from the PubChem (https://pubchem.ncbi.nlm.nih.gov/) database (Kim et al., 2019) in .sdf format and converted into .pdb structure using OpenBabel (https://openbabel.org/) (O’Boyle et al., 2011). The ligands were further energy minimized using Chem3D (ChemOffice 2002) MM2 energy minimization program.

For the receptor preparation, the structure of Glutathione Reductase (PDB ID: 3DK9) was downloaded from RCSB Protein Data Bank (rcsb.org). The receptor preparation was performed by deleting unwanted residues and water molecules and further subjecting the receptor to the energy minimization process using Chimera (Petterson et al., 2004). Ligand - Receptor visualization was done using LigPlot (Laskowski and Swindells 2011).

### Docking Protocol Validation

We docked Flavin Adenine Dinucleotide (FAD) with the GR crystal structure, using PatchDock server (https://bioinfo3d.cs.tau.ac.il/PatchDock/php.php; Duhovny et al., 2002; Schneidman-Duhovny et al., 2005), in order to validate the docking methodology, and we then compared the interacting residues of GR with the docked FAD molecules to the interacting residues of GR with FAD in the crystal structure (PDB). The RMSD between the bound FAD molecule and the crystal structure was determined to verify the validation.

### Molecular Docking

PatchDock (https://bioinfo3d.cs.tau.ac.il/PatchDock/php.php) online server (Duhovny et al., 2002; Schneidman-Duhovny et al., 2005) was employed for the molecular docking purpose. The active site of the receptor served as the binding site for ligands. The clustering RMSD values were set to 1.5 Å as suggested by the server for protein-ligand docking (Sangeet and Khan 2020; Sangeet et al., 2022a). The protein structure was uploaded in the receptor option and the phytochemicals were uploaded in the ligand option. To analyze the result, the PatchDock score and the corresponding Atomic Contact Energy (ACE) values were tabulated. As PatchDock gives an ensemble of possible structures, top 20 docked structures were downloaded and a rigorous selection procedure was followed based on the PatchDock score, ACE values and visualizing the docked structures to select the top hit phytochemicals. The visualization was carried out using LigPlot software (Laskowski and Swindells 2011).

### Molecular Dynamic Simulation

Molecular Dynamic Simulations were carried out in order to understand the dynamical behavior of the phytochemicals while interacting with Glutathione Reductase. The MD Simulations were carried out using GROMACS version 2018.1 (Abraham et al., 2015, Hess et al., 2008, Van Der Spoel et al., 2005) for 150ns. The protein-ligand preparation was followed before proceeding the simulations. The system was prepared using the CHARMM36 force field and the TIP3P water system. CGenFF (CHARMM General Force Field) (Vanommeslaeghe et al., 2010) was used to generate the ligand topology. The system was then solvated in a cubic box followed by energy minimization. Under NVT ensemble, the system’s temperature was gradually stabilized from 0K to 300K for 5 ns. The system was then put through an NPT ensemble simulation at 300K and 1.0 bar of pressure. The system coordinates were stored every 20ps during the 150ns production run for later post-processing analysis (Sangeet et al., 2022b).

### Root Mean Square Deviation (RMSD)

During the 150 ns simulation period, the structural and dynamic characteristics of the protein-ligand complexes were examined as the backbone RMSDs. The RMSD was calculated using the following equation as the average distance between the backbone atoms of the protein-ligand structures:

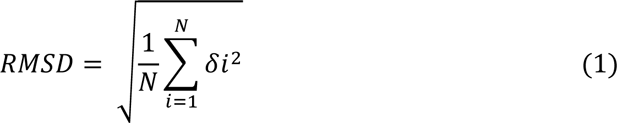

Where N is the total number of atoms taken into account for the computation and δ is the separation between each pair of equivalent atoms in N.

### Root Mean Square Fluctuation (RMSF)

The flexibility of each residue in the ligand-protein complexes was evaluated and depicted using the root mean square fluctuations (RMSF). The minimal fluctuation for all of the complexes was indicated by the protein ligand complex’s RMSF.

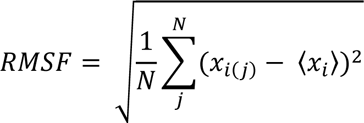

Where, 𝑥_𝑖𝑗_ denotes the coordinates (position) of the i^th^ Cα atom in the structure of the j^th^ model and 〈𝑥_𝑖_〉 denotes the averaged position of the i^th^ Cα atom.

### Radius of Gyration (RoG)

Calculations were made to determine the RoG for each complex backbone. Protein compactness is determined by the radius of gyration of the protein. A protein’s corresponding radius of gyration will have a reasonably constant value of RoG if it is folded consistently. On the other hand, the protein’s radius of gyration will be reflected by a fluctuating value of RoG if it is unstable or begins to unfold.

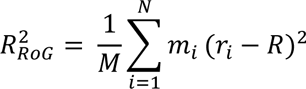

Where 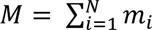 is the total mass and 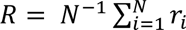 is the centre of mass of the protein consisting of N atoms.

### Solvent Accessible Surface Area (SASA)

The surface area of a biomolecule that is reachable by a solvent is known as the accessible surface area (ASA) or SASA. Square angstroms are commonly used to characterize SASA measurement units. The SASA was calculated for protein as well as the complex formed by the interaction of best hit ligands with the protein. The detailed methodology of SASA is given in the supplementary file.

### Hydrogen Bond Analysis

Hydrogen bond interaction was calculated between the ligands and the protein to understand the interaction evolution of the ligand with the respective proteins. For the purpose of conducting an H-bond analysis, atoms were chosen to be possible donors and acceptors. Atom names, atom kinds, or other criteria specified in the simulation system was used to make this decision. Most of the time, the donor atom contains the hydrogen atom, whereas the acceptor atom is the heavy atom that interacts with the hydrogen. A distance criterion was used to determine if the donor and acceptor atoms are within a given cutoff distance after the donor and acceptor atoms were identified. The expected range of H-bond distances, which is frequently between 3Å and 4Å, was typically used to determine this cutoff value. Atoms that were close to this cutoff point were thought to be possible H-bond partners. An angle criterion was also applied in addition to the distance requirement to evaluate the linearity of the hydrogen bond. The angle that was produced between the donor, hydrogen, and acceptor atoms was measured using this criterion. An angle cutoff of between 120 and 180 degrees was typically used to determine a legitimate H-bond. The hydrogen bond evolution was calculated for 150 ns.

### Dynamic Cross – Correlation Matrix (DCCM)

Dynamic cross-correlation analysis was used to investigate the association between the protein residues. The input for the computation of the cross correlation was taken from the protein’s crystal structure (PDB ID: 3DK9). An output trajectory with a lot of structural ensembles was produced using the trajectory file that was received following the MD simulation of the protein with the top hit ligands. R package, Bio3D was used for dynamic cross-correlation analysis on the trajectory (Grant et al., 2021). This package outputs a matrix of all atom-wise correlations, 𝐶_𝑖,𝑗_, whose elements can be seen graphically as DCCM.

### Drug Likeness of top hit phytochemicals

Lipinski’s Rule of Five (RO5) is a set of variables that enables early screening of phytochemicals. By determining their Lipinski value, the phytochemicals were also tested against the pharmaceuticals on the market using the SWISS-ADME server (http://www.swissadme.ch/) (Daina et al., 2017).

### ADME-Tox studies

With the help of the ProTox-II (https://www.tox-new.charite.de/protox_II/; Banerjee et al., 2018)

and SWISS-ADME servers (https://www.swissadme.ch/) (Daina et al., 2017), adsorption, digestion, metabolism, excretion, and toxicity analysis were carried out. Drug similarity parameters, such as lipophilicity, solubility, and drug likeness score, were determined based on the uploaded structures of pharmaceuticals and phytochemicals.

## Results and Discussion

Inhibition of Glutathione Reductase (GR) holds significant ramifications across diverse disease contexts and has garnered attention as a promising target for the development of anticancer and antimalarial therapeutics (Schirmir et al., 1995; Davioud-Charvet et al., 2001; Biot et al., 2003; Fidock et al., 2004; Bauer et al., 2006). The pursuit of GR inhibition can be realized through various chemical agents. However, due to concerns pertaining to chemical-induced toxicity, the imperative arises to devise innovative strategies for precise GR targeting

*Houttuynia cordata*, recognized for its potential anticancer attributes (Kim et al., 2017), presents an appealing avenue, owing to its status as a naturally occurring repository of phytochemicals that exhibit diminished toxicity in comparison to synthetic counterparts. Leveraging the repertoire of phytochemicals inherent in *Houttuynia cordata*, we embarked upon computational endeavors against the GR enzyme, elucidating potential phytomolecules endowed with the capacity to impede GR functionality.

To corroborate the docking outcomes, a comprehensive assessment ensued, involving 150 ns molecular dynamics simulations to unravel the dynamic interplay between the identified phytocompounds and the GR enzyme. Furthermore, an evaluation of Absorption, Distribution, Metabolism, Excretion, and Toxicity (ADME-Tox) properties was undertaken, affording insights into the pharmacological attributes of the foremost phytocompound candidates.

### Active Site Identification

The Richards’ Solvent Accessible Surface Area/Volume (SASA) was calculated to be 1641.189 Å^2^ and 1606.932 Å^3^ respectively (Fig. 1b). The calculated SASA was confirmed by analyzing the interaction of FAD in the co-crystallized form with GR (Fib. 1a). The interaction of FAD with GR in the co-crystallized form confirmed the authenticity of the computed active site.

**Fig. 1:**
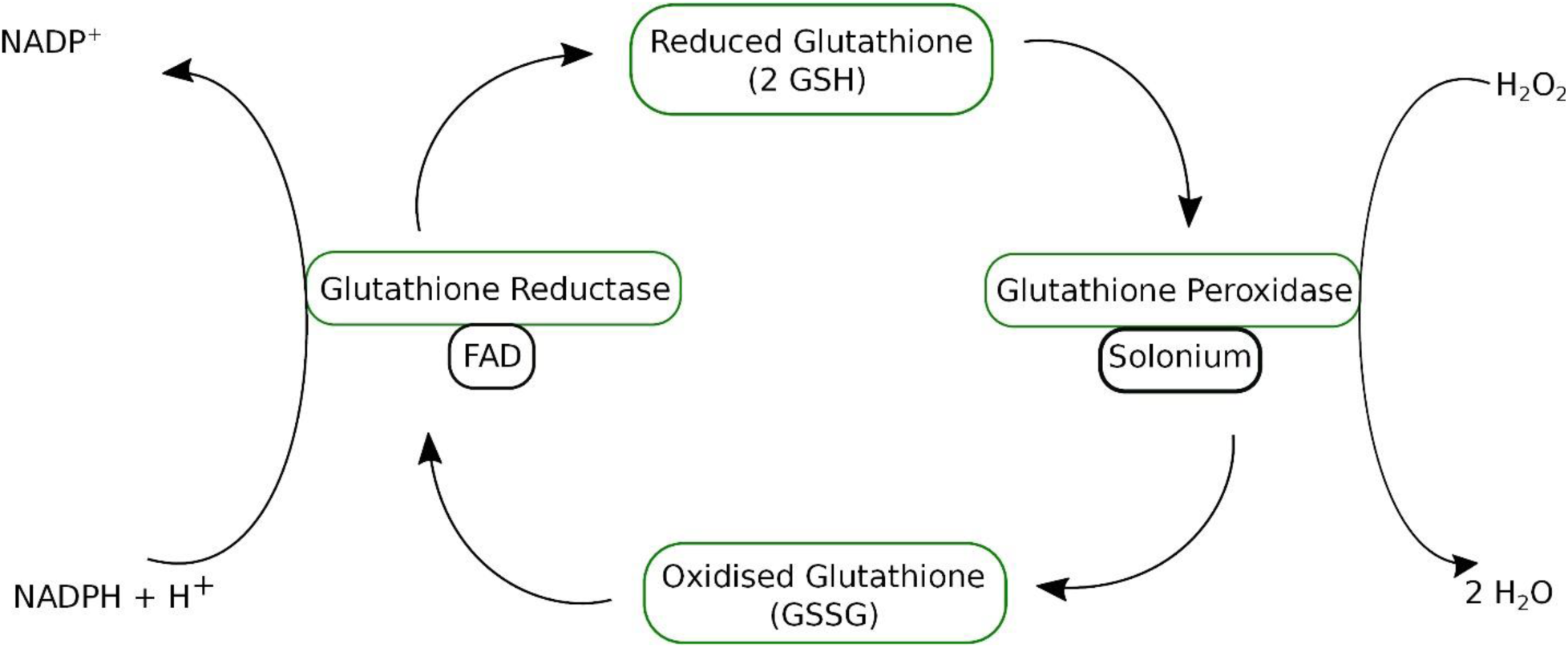
Glutathione Reductase Cycle. Glutathione Peroxidase converts H_2_O_2_ to 2H_2_O using reduced Glutathione (GSH) and Glutathione Reductase converts NADPH + H^+^ into NADP^+^ using oxidised Glutathione (GSSG)

### Phytochemical Data Collection

Dr Duke’s Phytochemical and Ethnobotanical Database (Duke 2016) was used to find the phytochemicals of *Houttuynia cordata* that display the antioxidant properties. A total of 13 phytochemicals were selected for the molecular docking studies. The top hit phytochemicals are represented in Fig.3. The other relevant phytochemical detail is presented in Table S1 (see Supplementary Table 1).

### Ligand and Receptor Preparation

The selected phytochemicals and the receptor were prepared by deleting any unwanted residues and water molecules that might potentially interfere with the docking process. Further, both the ligands and the receptor were energy minimized using Chem3D and protein was energy minimized using Chimera (UCSF Chimera 2004). The crystal structure of the protein is shown in Fig. 2a. We carried out a thorough docking investigation including the interaction between Flavin Adenine Dinucleotide (FAD; Fig. 4a) and the crystalline architecture of Glutathione Reductase (GR) in order to validate the applied docking procedure. The amino acid residues involved in interactions with FAD molecules that were discovered by docking (Fig. 4) versus those residues mediating interactions between GR and FAD within the crystallographic structure (PDB) were compared meticulously. Our research showed that the docked FAD had a strong binding affinity to the catalytic site of GR, which was supported by a high PatchDock score of 4986 and a particularly high binding energy of −195.76 kcal/mol. The three-dimensional location of the bound FAD was then aligned with the ligand in the native crystal structure using a superimposition procedure (Fig. 4b). When this alignment was evaluated, the RMSD (Root Mean Square Deviation) difference was 0.57Å. This outcome concretely substantiated the congruence of our applied docking protocol.

**Fig. 2:**
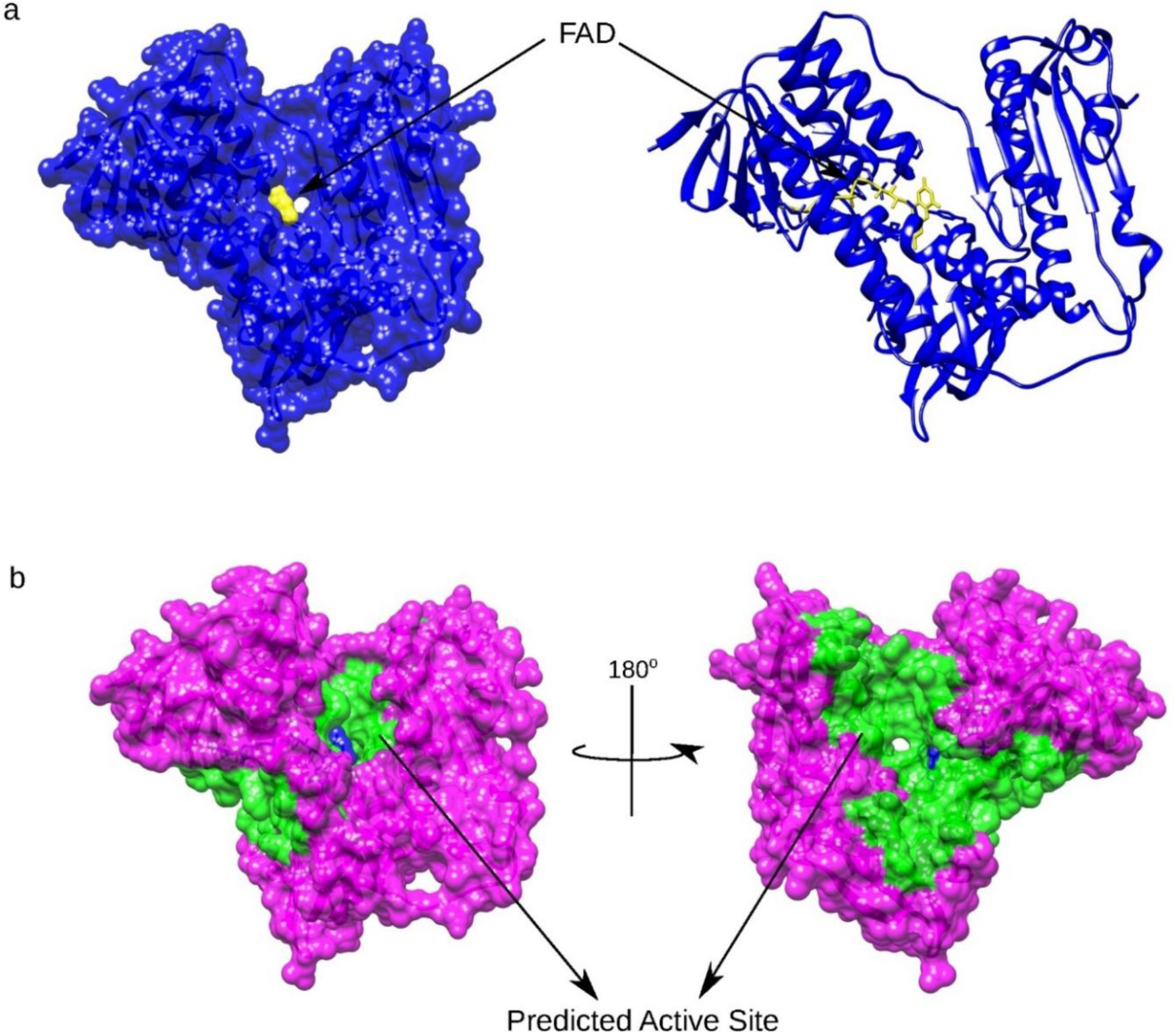
Structure of Human Glutathione Reductase (PDB ID: 3DK9) (a) Surface view and crystal structure of human glutathione reductase (blue) in association with the co-crystallized ligand Flavin-Adenine Dinucleotide (FAD, yellow). (b) Binding pocket (green) of human glutathione reductase (magenta) as predicted by CASTp server.

**Fig. 3:**
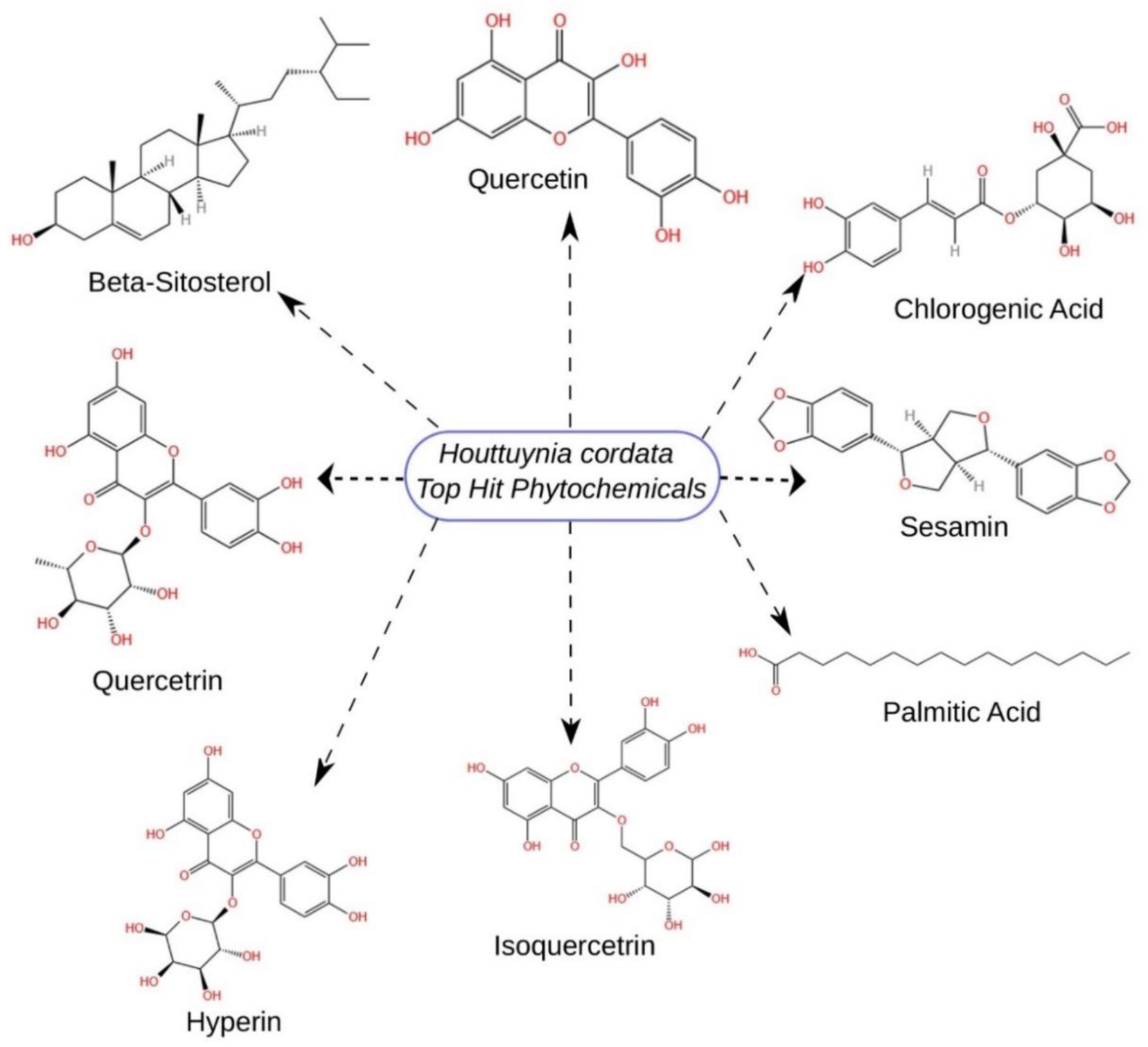
2D structure of the top phytochemicals of *Houttuynia cordata*. Out of the top eight phytochemicals, Quercetin, Quercitrin and Sesamin demonstrated a better binding affinity with the human glutathione reductase.

**Fig. 4:**
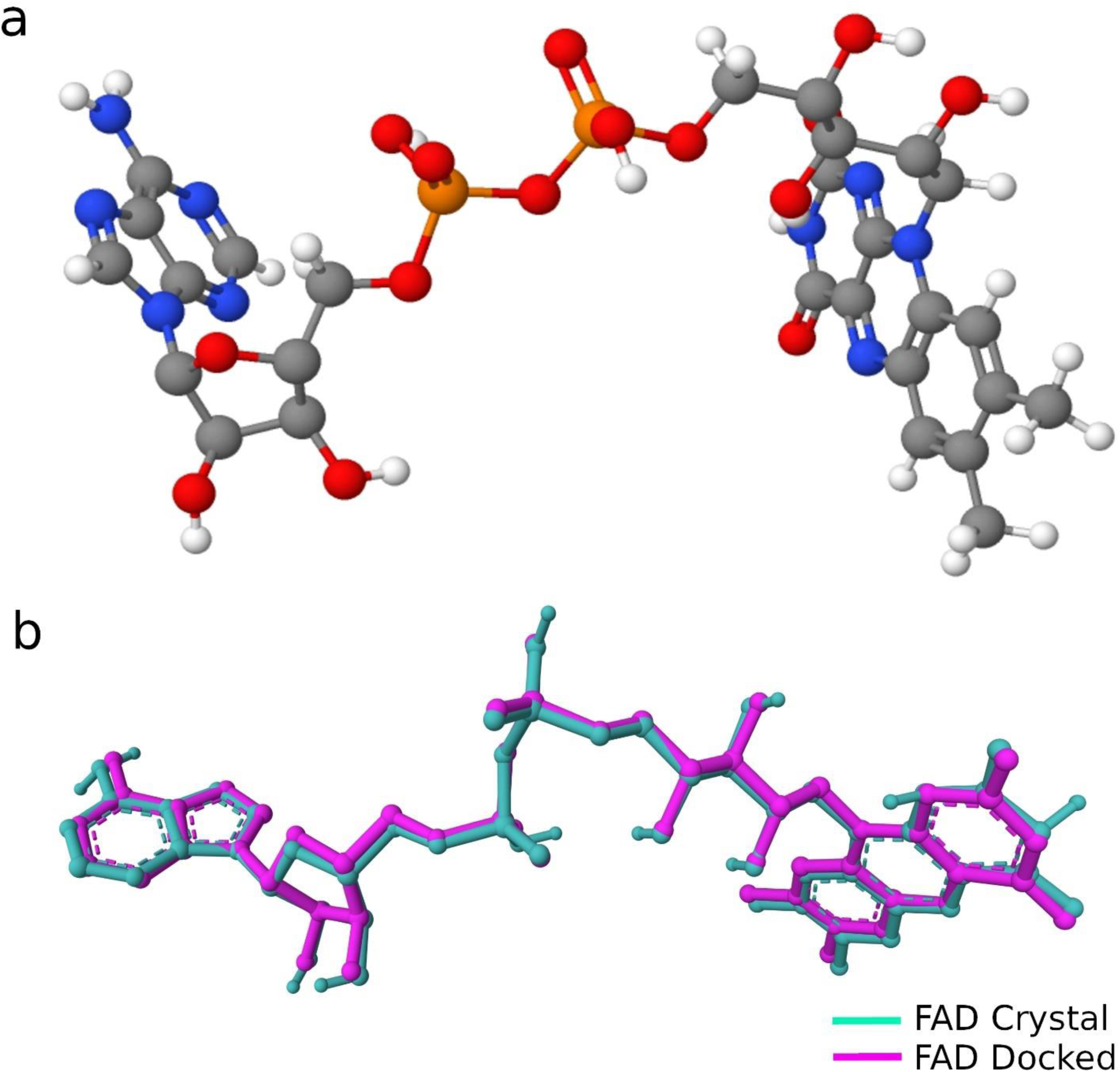
Flavin Adenine Dinucleotide (FAD) analysis showing the (a) 3D structure of FAD and (b) Superimposition of FAD in crystal (cyan) with the docked FAD (magenta).

On analyzing the interaction of the co-crystallized ligand (Flavin-Adenine Dinucleotide, FAD) with the protein, we identified that FAD forms hydrogen bonding with Glu50, Ala130, Ser51, Thr57, Lys66, Thr339, Asp331, Gly31 and Ser30. Apart from showing an extensive hydrogen bonding, it also interacts with other residues via hydrophobic interactions as represented in Fig. 5.

**Fig. 5:**
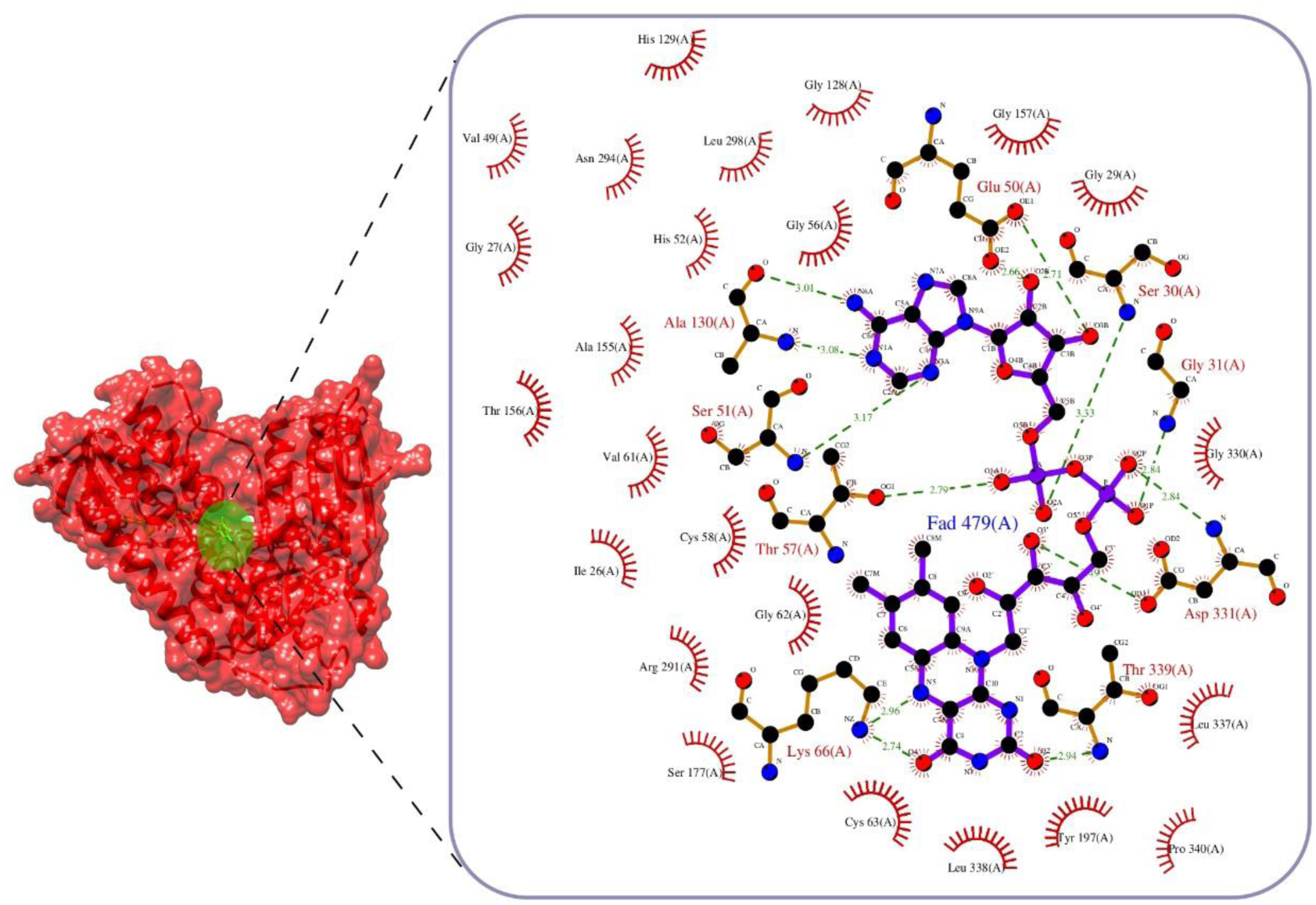
Molecular Interaction of Flavin Adenine Dinucleotide with Human Glutathione Reductase. Surface view of GR (red) showing the position of FAD (yellow). Residual interaction between FAD and GR (box) showing extensive amount of Hydrogen and Hydrophobic interaction.

### Molecular Docking

PatchDock online server (https://bioinfo3d.cs.tau.ac.il/PatchDock/php.php) was utilized for performing molecular docking studies between the ligands and the receptor. The active site of the receptor served as the binding site for the ligands. The binding affinity between ligands and the receptor was ascertained through the PatchDock Score (PS) and the Atomic Contact Energy (ACE) value (Sangeet and Khan 2020). We discovered eight phytochemicals (Fig. 3) that bind to the protein’s active site after performing molecular docking between the phytochemicals and the protein. Three of these eight phytochemicals—Quercetin, Quercitrin, and Sesamin—showed a greater affinity for binding than the others. Along with hydrophobic interactions with Thr57, Asp331 and Ser30, Quercetin displayed hydrogen bonding with Lys66 and Thr339 as well with a PatchDock score of 4538 and an ACE value of −180.3 kcal/mol (Fig. 6a). Quercitrin on the other hand, showed a hydrogen bond interaction with Leu337, Leu338, Cys63, Thr57 and Ser177 with a PatchDock score of 5530 and an ACE value of −220.96 kcal/mol. Moreover, it exhibited hydrophobic interaction with Lys66, Thr339 and Asp331 (Fig. 6b). Furthermore, Sesamin showed hydrogen bond interactions with Lys66 and Ile198 with a PatchDock score of 5484 and an ACE value of −243.19 kcal/mol. Sesamin also formed hydrophobic interactions with Thr57, Thr339, Asp331 and Ser30 (Fig. 6c). After examining the interactions of the top three phytochemicals, we discovered that they exhibited superior binding behavior than other phytochemicals that had been screened, demonstrating a higher binding affinity for the protein’s active region. Table 1 shows the interacting residues for the most popular hit phytochemicals. Details of the protein’s interactions with additional well-known phytochemicals are included in the Supplementary Information (see Supplementary Table S2).

**Fig. 6:**
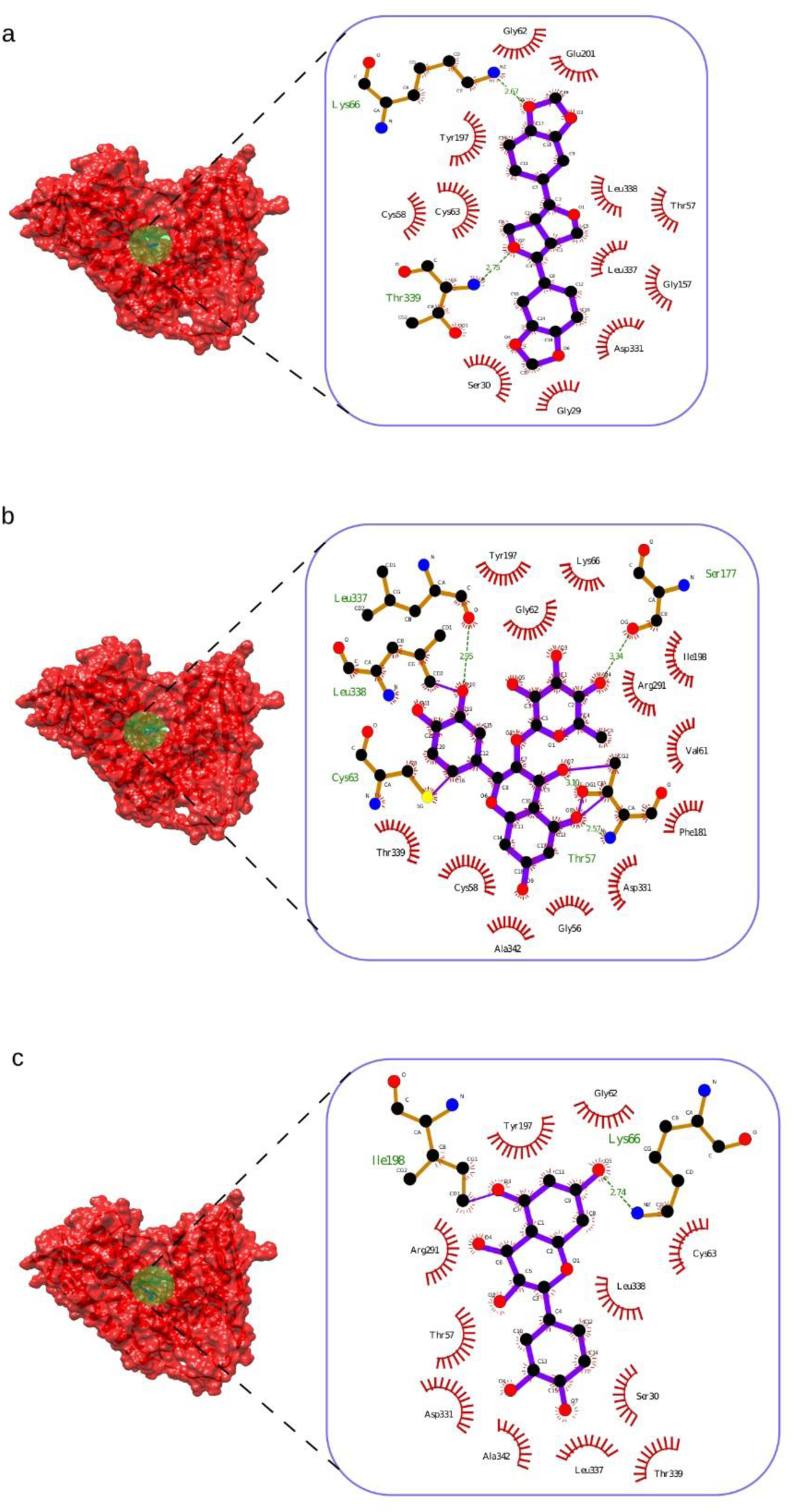
Molecular Interaction of top hit phytochemicals with GR. (a) Quercetin exhibits a hydrogen bond interaction with Lys66 and Thr339 (b) Quercitrin shows hydrogen bonding with Leu337, Leu338, Cys63, Thr57 and Ser177 (c) Sesamin shows hydrogen bond interaction with Lys66 and Ile198

**Table 1:** Molecular information about FAD and the top hit phytochemicals interacting with GR.

| Phytochemical | PatchDock<br>Score | Atomic Contact<br>Energy (kcal/mol) | Interacting Residues |
| --- | --- | --- | --- |
| FAD | 4986 | -195.76 | His129, Val49, Asn294, Leu298, Gly128, Gly27, His52, Gly56, Ala155, Thr156, **Ala130\***, **Ser51\***, Val61, Ile26, Cys58, **Thr57\***, Gly62, Arg291, Ser177, **Lys66\***, Cys63, Leu338, Tyr197, Pro340, Leu337, **Thr339\***, **Asp331\***, Gly330, **Gly31\***, **Ser30\***, Gly29, **Glu50\***, Gly157 |
| Quercetin | 4538 | -180.3 | **Lys66\***, Gly62, Glu201, Leu338, Thr57, Leu337, Gly157, Asp331, Gly29, Ser30, **Thr339\***, Cys58, Cys63, Tyr197 |
| Quercitrin | 5530 | -220.96 | **Leu337\***, Tyr197, Lys66, Gly62, **Ser177\***, Ile198, Arg291, Val61, Phe181, Asp331, **Thr57\***, Gly56, Ala342, Cys58, Thr339, **Cys63\***, **Leu338\*** |
| Sesamin | 5484 | -243.19 | **Ile198\***, Tyr197, Gly62, **Lys66\***, Cys63, Leu338, Ser30, Thr339, Leu337, Ala342, Asp331, Thr57, Arg291 |

*\*Residues showing Hydrogen Bonding with GR*

### Molecular Dynamic Simulation

A 150ns MD simulation was performed on the protein-ligand complexes’ molecular simulations. The top phytochemical interactions with the protein were computed using the relevant RMSD, RMSF, Hydrogen Bond, Solvent Accessible Surface Area and Radius of Gyration values. The ligand conformations corresponding to the simulation times of 30 ns, 90 ns, and 150 ns were retrieved for respective ligands, FAD (Fig. S1 a/b/c), Quercetin (Fig. S1 d/e/f), Quercitrin (Fig. S1 g/h/i), and Sesamin (Fig. S1 j/k/l). These configurations demonstrate how the ligand evolved throughout the simulation process, showing that it explored different configurational options in order to find the best stable configuration that complemented the protein’s binding pocket. We further analyzed the Ramachandran plot of Glutathione Reductase with the respective ligands for 30 ns, 90 ns and 150 ns (Fig. S2) and observed that the respective ligands didn’t affect the structural integrity of the protein suggesting their good binding profile.

### Root Mean Square Deviation (RMSD)

The stability of the protein-ligand combination throughout the course of an MD simulation is shown by the term “Root Mean Square Deviation” (RMSD). Throughout the 150ns MD Simulation, Quercetin, Quercitrin, and Sesamin remained steady. The RMSD fluctuations for Quercetin, Quercitrin, and Sesamin, respectively, were within the permissible limits of 0.15-0.4 nm, 0.1-0.35 nm, and 0.2-0.25 nm (Fig. 7a). Compared to the most popular phytochemicals, FAD showed a somewhat similar variability within 0.1–0.35 nm (Fig. 7a black). Quercetin demonstrated an increase in the RMSD till 50ns beyond which it suddenly peaked its value at 60 ns beyond which it stabilizes throughout the simulation with an average RMSD value of 0.31 nm (Fig. 7a red). Quercitrin showed a sudden peak in the value of RMSD at 23 ns after which the behavior of the binding between Quercitrin and protein assumed a stable trajectory (Fig. 7a green). Throughout the trajectory, Sesamin demonstrated a consistent binding with an average RMSD range of 0.2-0.25nm (Fig. 7a blue). Additionally, the constrained confirmational changes during the simulation process are confirmed by the histogram of RMSD of various ligands (Fig. 7b). The RMSD distribution of FAD shows a bimodal distribution with a peak at 0.28 nm. A similar bimodal distribution is obtained for Sesamin where the peak is obtained at 0.2 nm. This bimodal behavior suggests that the binding of the respective ligand makes the protein take two distinct conformations throughout the MD simulation. On the other hand, the RMSD distribution of Quercetin shows a broad distribution suggesting that after the binding event, protein assumes an ensemble of structures. Moreover, the RMSD distribution of Quercitrin depicts some minor humps in the distribution at 0.15 nm and 0.19 nm respectively. But a distinct peak is obtained at 0.24 nm suggesting that the binding behavior of Quercitrin makes the protein take a distinct conformation. This is analysis is supported when we analyze the structural evolution of Quercitrin (Fig. S1 g/h/i). It can be seen that Quercitrin acquires minor side chain changes but not a distinct structural change which maybe the cause of getting the small distribution peaks (Fig. 7b). As is evident from the histogram plot, the confirmational fluctuation for FAD and Quercetin is very broad, resembling that the ligand fluctuates more to explore the conformational space (Fig. 7b black and red respectively). This is confirmed by the structural evolution of FAD (Fig. S1 a/b/c) and Quercetin (Fig. S1 d/e/f) throughout the simulation trajectory.

**Fig. 7:**
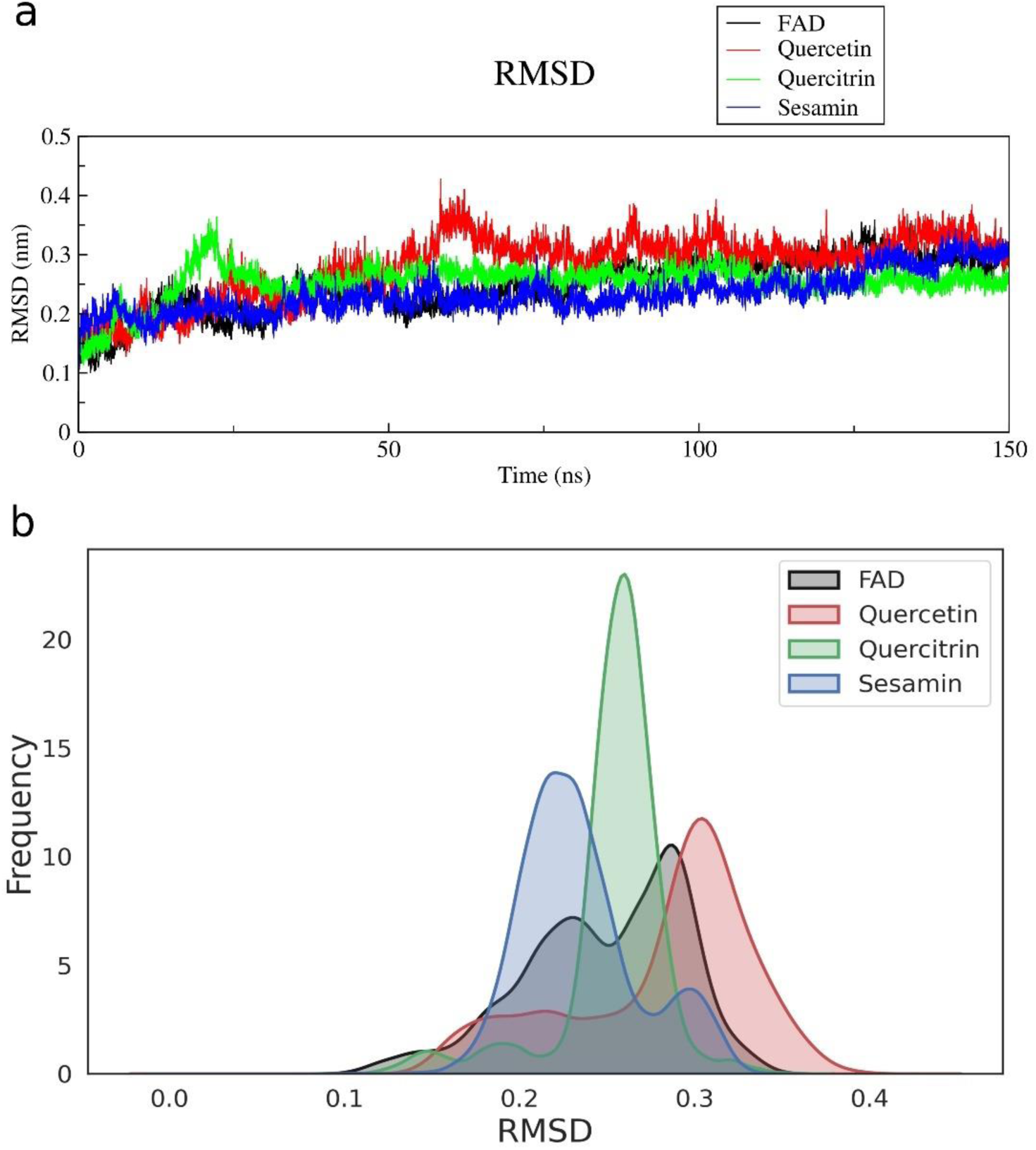
Root Mean Square Deviation (RMSD) of the top hit phytochemicals for a molecular simulation of 150ns. (a) RMSD trajectory of the top three phytochemicals along with the co-crystallized ligand, FAD. The trajectory reveals that throughout the simulation the binding profile of all the top hit phytochemicals demonstrated a stable binding with fluctuation lying in the permissible limit. (b) Histogram plot of RMSD of the top hit phytochemicals and FAD. The histogram shows that for Quercitrin the RMSD fluctuation was very narrow with majority of the fluctuation occurring in the range of 0.25 nm. Sesamin and FAD had a bimodal distribution in the histogram plot suggesting that the ligands make the protein acquire two distinct structural orientations throughout the simulation trajectory.

### Root Mean Square Fluctuation (RMSF)

At the residual level, the Root Mean Square Fluctuation (RMSF) shows flexibility (Sangeet et al., 2021). All of the ligands showed very little variation. The steady behavior of the functionally significant residues between the corresponding ligands and the protein is shown by the RMSF fluctuation plots (Fig 8a). It was found that Quercitrin showed a lower fluctuation as compared to other phytochemicals (Fig. 8a green). Almost all the top hit phytochemicals resembled a similar behavior, there were regions where Sesamin and Quercitrin showed a better stability than Quercetin. Moreover, we characterized the RMSF in three distinct regions corresponding to the active site of the protein (Fig. S3). It can be seen that in region 1 (R1) the RMSF of Quercitrin and Sesamin were lower as compared to Quercetin and FAD suggesting that the binding of Quercitrin and Sesamin lowers the fluctuation in the active site of the protein. The fluctuation corresponding to residues 26-31, 49-58 and 61-66 constitute region 1. As we move ahead in residue numbers, we can observe the higher fluctuation of FAD and Quercetin corresponding to a value of 0.4nm (Fig. 8a black and red). Same pattern can be observed in region 2 (R2) and region 3 (R3) as well. Moreover, in the overall trajectory, some unique behavior was also observed. It was found that the binding of all the phytochemicals induce a higher fluctuation in the distant regions from the active site, especially in the region post 400 residue number (Fig. 8a). Furthermore, we identified three distinct regions in the protein that showed a higher fluctuation whenever a ligand binds to the Glutathione Reductase. We characterize these sites as Fluctuating Zone 1 (residue 90-95), Fluctuating Zone 2 (268-274) and Fluctuating Zone 3 (405-415) (Fig. 8a).

**Fig. 8:**
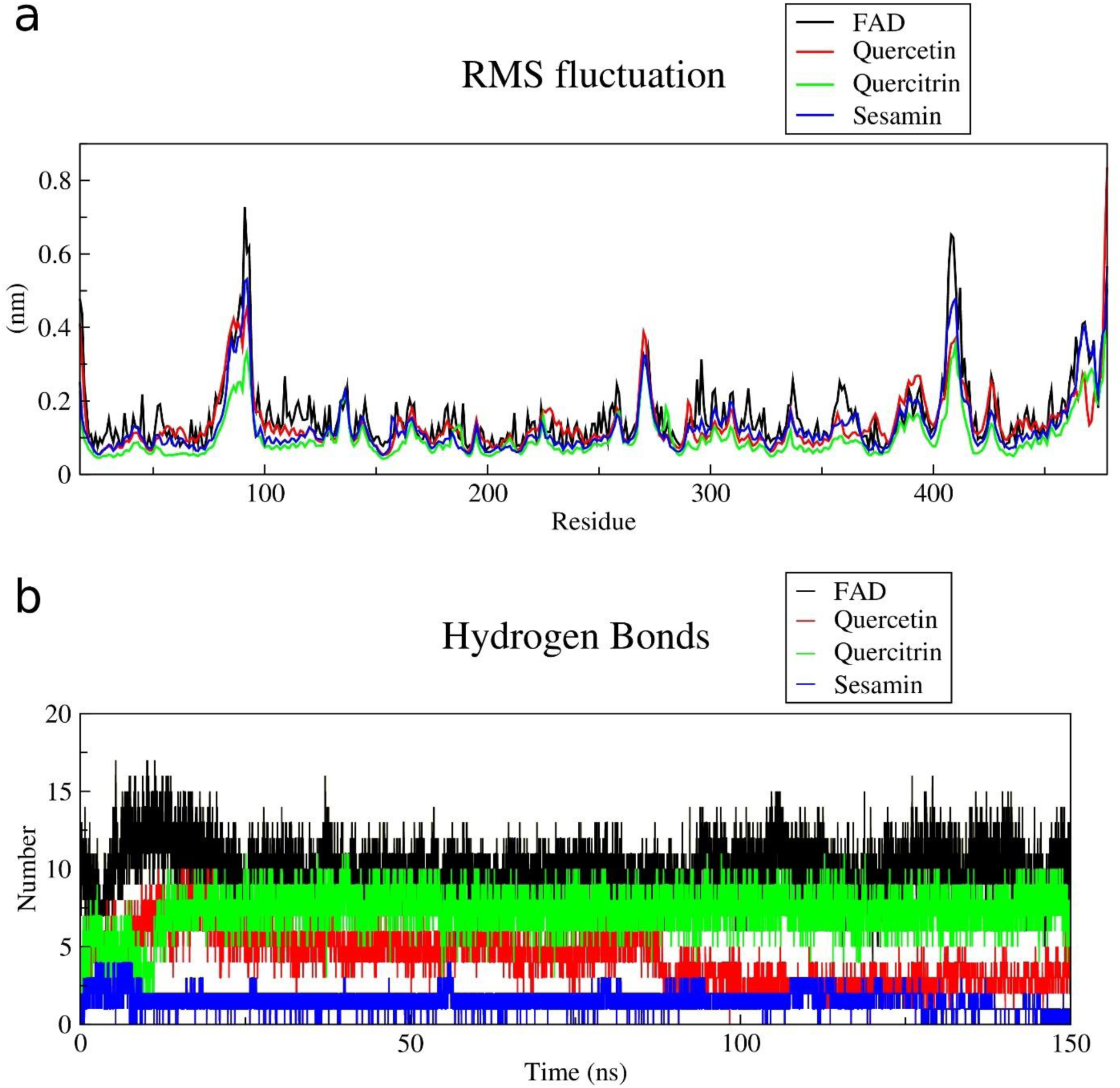
Root Mean Square Fluctuation (RMSF) and Hydrogen Bond analysis of Glutathione Reductase and the complex in conjugation with top hit phytochemicals. (a) RMSF curve corresponding to the binding of FAD (black), Quercetin (red), Quercitrin (green) and Sesamin (blue) (b) Hydrogen Bond analysis of GR when top hit phytochemicals interact with the protein.

### Hydrogen Bond

The hydrogen bond interaction between the protein and ligand was analyzed to understand the plausible mode of binding. It was found that the interaction of FAD with GR formed a higher number of hydrogen bonds (n=16) during the initial simulation and then slowly stabilizes with 12 hydrogen bonds on average post 25 ns (Fig. 8b black). Quercetin also showed a similar hydrogen bond pattern where it forms around 10 hydrogen bonds during the initial simulation and then slowly stabilizing at 6 hydrogen bonds on average post 25 ns (Fig. 8b red). It was interesting to see that Quercetin maintained 6 hydrogen bonds on average till 75 ns and then suddenly dropping down to 3 hydrogen bonds on average through the rest of the simulation. Quercitrin also resembled a similar behavior with the number of hydrogen bonds varying between 2-10 (Fig. 8b green). Quercitrin also resembled a unique behavior where the number of hydrogen bonds were fluctuating initially between 0-10ns after which it stabilized to a range of 6-10 hydrogen bonds with the maximum number going to 11 hydrogen bonds. This behavior shows that the compound has strong binding affinity to the active site of GR. The initial fluctuating hydrogen bonds might indicate an exploratory binding process where the compound establishes various interactions with the protein eventually obtaining a stabilization in hydrogen bonds indicating the specific interactions that might be dominant in nature. Sesamin, on the other hand, showed a stabilized hydrogen bond number ranging between 1-3 throughout the simulation process with a maximum number of hydrogen bond corresponding to 4 (Fig. 8b blue).

### Radius of Gyration (RoG)

The Radius of Gyration is a symbol for the stability, compactness, and folding of proteins (Sangeet et al., 2022c). The outcomes imply that following the binding of FAD, Quercetin, Quercitrin, and Sesamin, the protein remained relatively compact and stable during the course of the 150 ns simulation (Fig. 9a). It was observed that the binding of FAD induces a RoG fluctuation in the range of 2.4-2.47 nm (Fig. 9a black). The fluctuation marginally goes down after 30ns depicting that the compactness of the protein increases after 30 ns but the RoG increases slightly beyond 125 ns depicting the unfolding of the protein structure. On the other hand, Sesamin showed a better behavior in terms of making the protein more compact as compared to other two phytochemicals (Fig. 9a blue). This behavior confirms the role of phytochemicals in creating a tensed structure for the protein. This behavior might be the result of limitation in structural movement due to the binding of the phytochemicals to the active site of the protein. Moreover, Quercitrin also resembled a similar RoG behavior as that of FAD and Sesamin indicating that the ligand is efficient in making the protein more compact (Fig. 9a green). On the other hand, Quercetin showed a higher RoG value as compared to other phytochemicals indicating that the binding phenomena of Quercetin makes the protein less compact (Fig. 9a red). This behavior is further confirmed by the distribution curve of RoG where the distribution curve of Quercetin has a peak value of 2.46 nm (Fig. 9b red). Whereas the distribution curve of FAD (Fig. 9b black) and Sesamin shows almost a similar peak with Sesamin showing a minor hump/shoulder region in the distribution curve (Fig. 9b blue). Quercitrin on the other hand shows a distinctive peak at a value of 2.40 nm (Fig. 9b green) suggesting that the binding of Quercitrin to the protein makes the protein compact and assume a stable configuration.

**Fig. 9:**
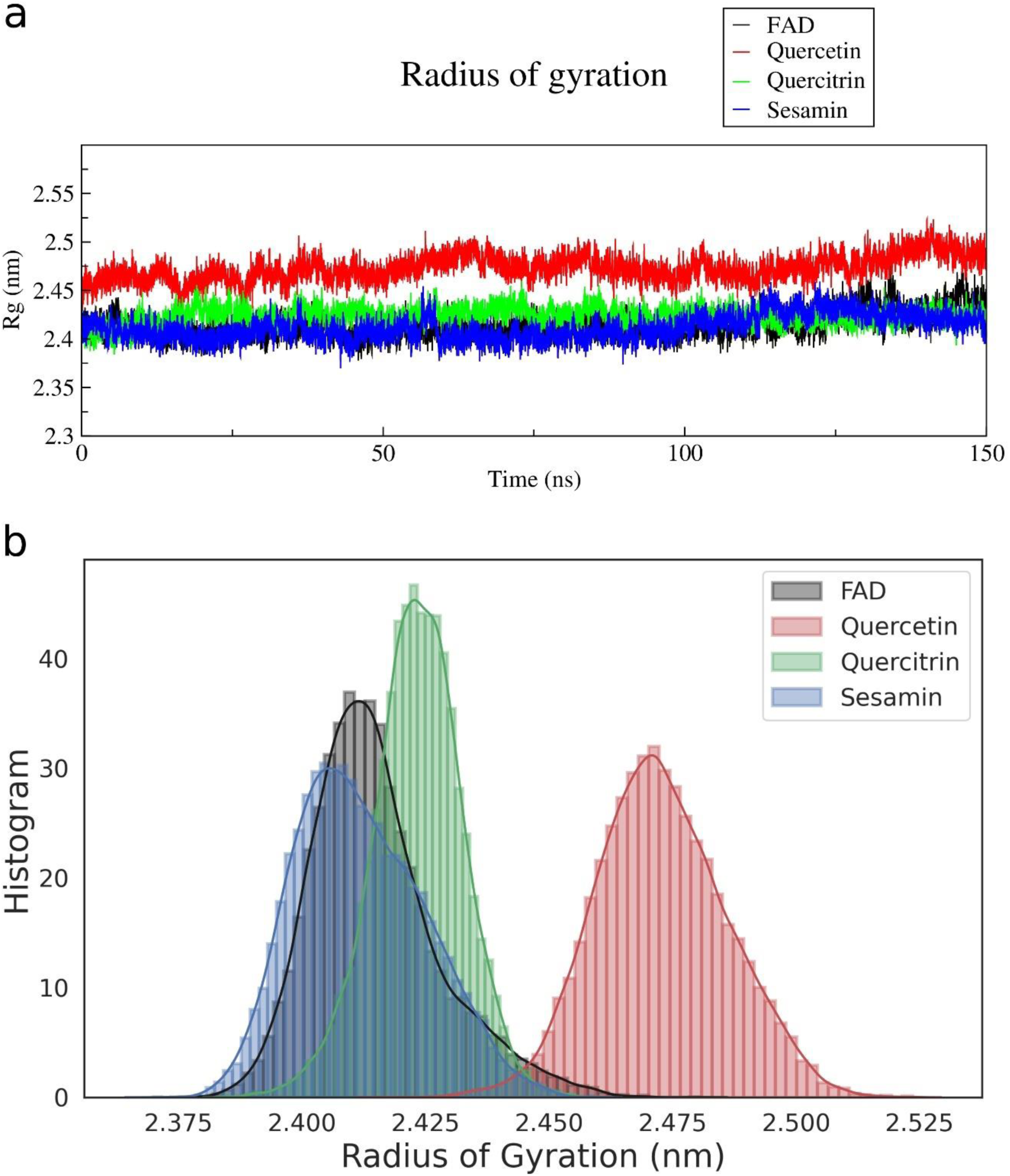
Radius of Gyration for the top hit phytochemicals with GR. (a) Radius of gyration of GR in the presence of FAD (black), Quercetin (red), Quercitrin (green) and Sesamin (blue) (b) Distribution of Radius of Gyration of the respective ligands with GR

### Solvent Accessible Surface Area (SASA)

SASA analysis captures the surface area of the protein which is available to the solvent where a higher SASA value represents a relative expansion and vice versa. As is seen in Fig. 10a, when GR interacts with FAD it starts with a SASA values of 230 nm^2^, reaches its highest value of 245 nm^2^ at around 25 ns and then gradually stabilizes at 245 nm^2^ at the end of the simulation (Fig. 10a black). Quercetin also follows a similar trend as that of FAD reaching its maximum value of 255 nm^2^ at 60 ns and then gradually fluctuating at a value of 245 nm^2^ the end of the trajectory (Fig. 10a red). Quercitrin, on the other hand, starts with a SASA value of 230 nm^2^ and reaches its maximum value of 250 nm^2^ at 20 ns, gradually fluctuating in the range of 240-245 nm^2^ and then again reaching the stable value of 245 nm^2^ at the end of the trajectory (Fig. 10a green). Sesamin started with a SASA value of 230 nm^2^ and reached its maximum value of 248 nm^2^ at 8 ns before it stabilized to 235 nm^2^ at the end of the trajectory (Fig. 10a blue). Moreover, from the distribution curve it can be seen that except for FAD all the three phytochemicals showed a unimodal distribution with a peak value of 232 nm^2^, 241 nm^2^ and 243 nm^2^ for Sesamin, Quercitrin and Quercetin respectively (Fig. 10b blue, green and red curve). FAD on the other hand showed a bimodal distribution that can be attributed due to its large size attaining a conformational change during the course of simulation. Because of structural change, more solvent molecules can access the surface of the active site. This is reflected as a bimodal distribution where the first peak for FAD (Fig. 10b black) can denote a structural configuration where the ligand is effectively interacting with the active site whereas the second peak, having a higher SASA value can correspond to a structural change which allows more solvent molecules to interact with the protein.

**Fig. 10:**
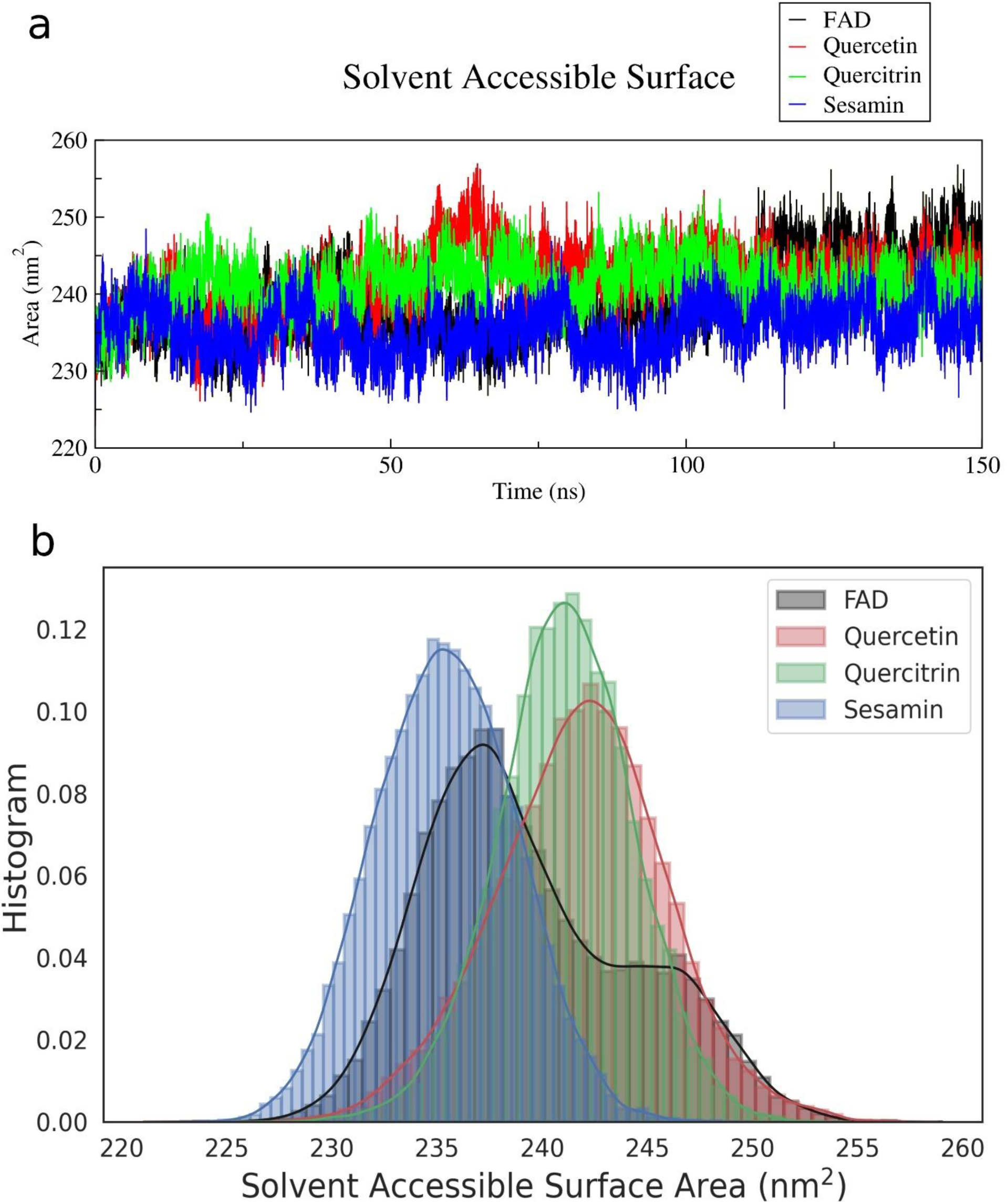
Solvent Accessible Surface Area (SASA) of top hit ligands with GR. (a) SASA of GR in the presence of FAD (black), Quercetin (red), Quercitrin (green) and Sesamin (blue) (b) Distribution of SASA of respective ligands with GR

### Dynamic Cross Correlation

To investigate the interplay between the protein and ligands – FAD, Quercetin, Quercitrin and Sesamin - dynamic cross-correlation matrix (DCCM) analysis was performed. The DCCM quantifies the correlations in motion between residues within a protein structure over time. Our analysis focused on understanding how the interactions with these ligands influence the dynamic behavior of the protein, with particular attention to the active site. To decipher the impact of the ligands on the protein’s active site, we focused on the region known to play a crucial role in substrate binding. By examining the DCCM entries corresponding to the active site residues, we gained insights into how each ligand modulates the correlated motions within this critical region. We observed that FAD binding induces anticorrelation in R2 and R3 regions (Fig. 11a yellow box) when compared to the R1 region. Although there were regions showing some minor positive correlation, but majority of the regions depicted a negative correlation suggesting that the binding of FAD to the protein induces a positive correlation in the R1 domain depicting a positively linked movement of one residue to the other. Quercetin also showed an anticorrelation in the R2 and R3 domain as compared to R1 domain (Fig. 11b yellow box). But compared to FAD, majority of the residues in this region showed no correlation suggesting an independent dynamic behavior of the residues. This phenomenon might suggest why quercetin had a similar dynamic behavior as that of FAD when bound to the protein but didn’t show quality behavior as compared to other two phytochemicals. Quercitrin, on the other hand, showed a comparative amount of positive and negative correlation in R2 domain and almost no correlation in R3 domain (Fig. 11c yellow box). This observation could point to the fact that when Quercitrin interacts with the protein, it induces a positive correlation which might be the reason why the dynamic behavior of quercitrin is better. Sesamin also resembled a similar behavior as that of Quercitrin showing a negative correlation in R2 domain and a mix of positive, negative and no correlation in R3 domain (Fig. 11d yellow box). But we also observed a unique behavior when the respective ligands bind to the protein. When FAD binds to the protein it induces a mix of negative and positive correlation in the region extending from residue 300-450 (Fig. 11a blue box). But it showed a positive correlation from 400-420. This behavior might suggest why in RMSF fluctuation, when FAD binds to the protein, the region 400-420 showed a highest RMSF fluctuation (Fig. 8a black). But the binding of Quercetin induces a negative correlation (Fig. 11b blue box) suggesting a dynamic behavior that reduces the fluctuation in the specific domain (Fig. 8a red). In the case of Quercitrin, it sowed a mix of negative and no correlation (Fig. 11c blue box) suggesting why the ligand is reducing the RMSF fluctuation in the region 400-420 whereas when Sesamin binds to the protein it induces a mix of positive and no correlation in the region 400-420 (Fig. 11d blue box). This analysis reveals why with this type of behavior Sesamin shows a higher fluctuation than Quercetin and Quercitrin (Fig. 8a blue).

**Fig. 11:**
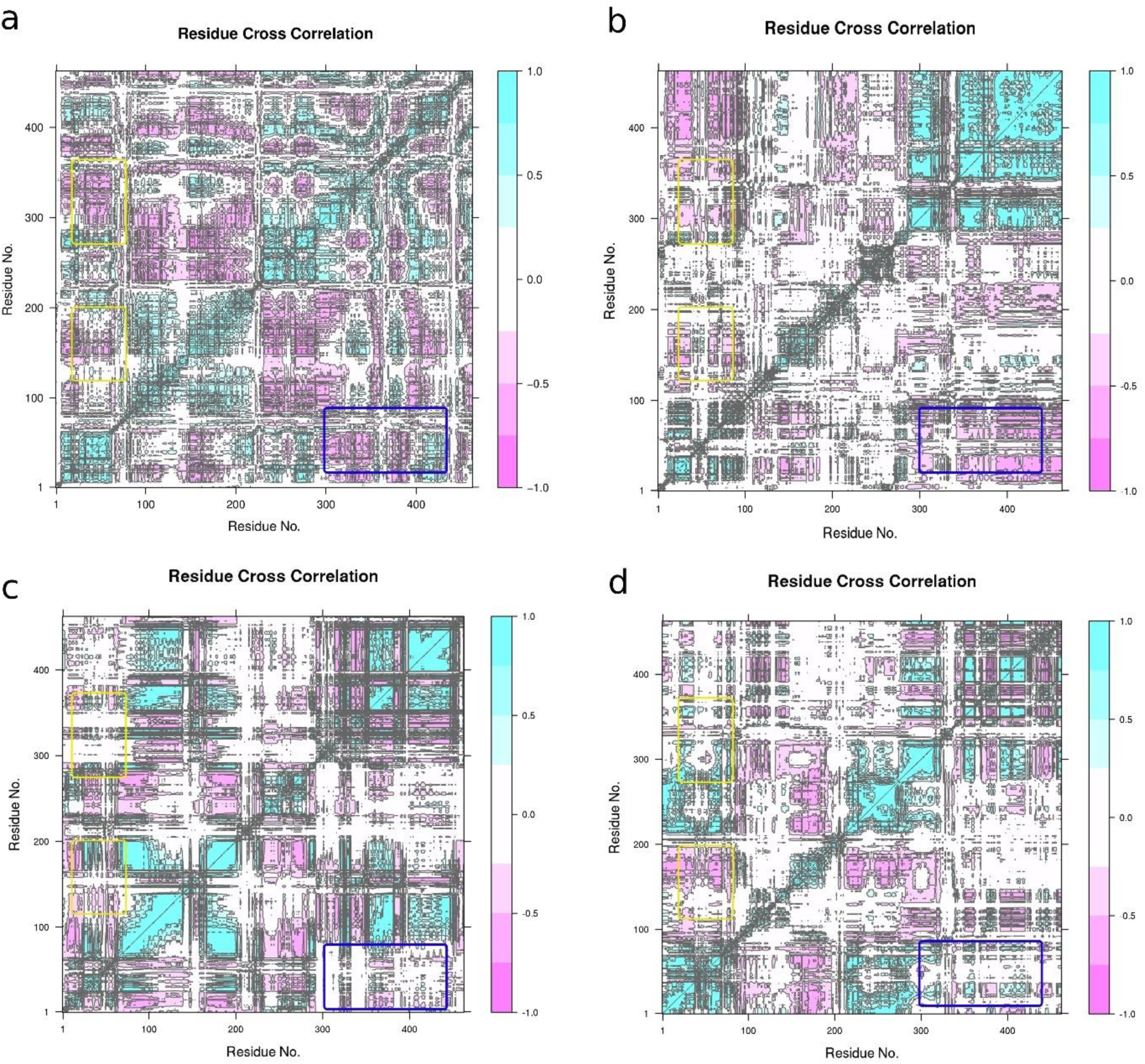
Dynamic Cross Correlation Matrix (DCCM) analysis of Glutathione Reductase in the presence of (a) FAD (b) Quercetin (c) Quercitrin and (d) Sesamin

### Drug Likeness of Phytochemicals

Drug likeness property of a compound characterizes them in terms of their potential to act as a drug. We employed Swiss-ADME server (https://www.swissadme.ch/; Daina et al., 2017) and Protox-II server (https://www.tox-new.charite.de/protox_II/; Banerjee et al., 2018) to calculate the Drug likeness and Toxicity properties of the top hit phytochemicals. Quercetin and Sesamin followed the Lipinski’s rule by showing no violation to the parameters. Quercitrin, on the other hand showed two violations by exceeding the limit of Hydrogen Bond Donors and Hydrogen Bond Acceptors (Table 2).

**Table 2:**
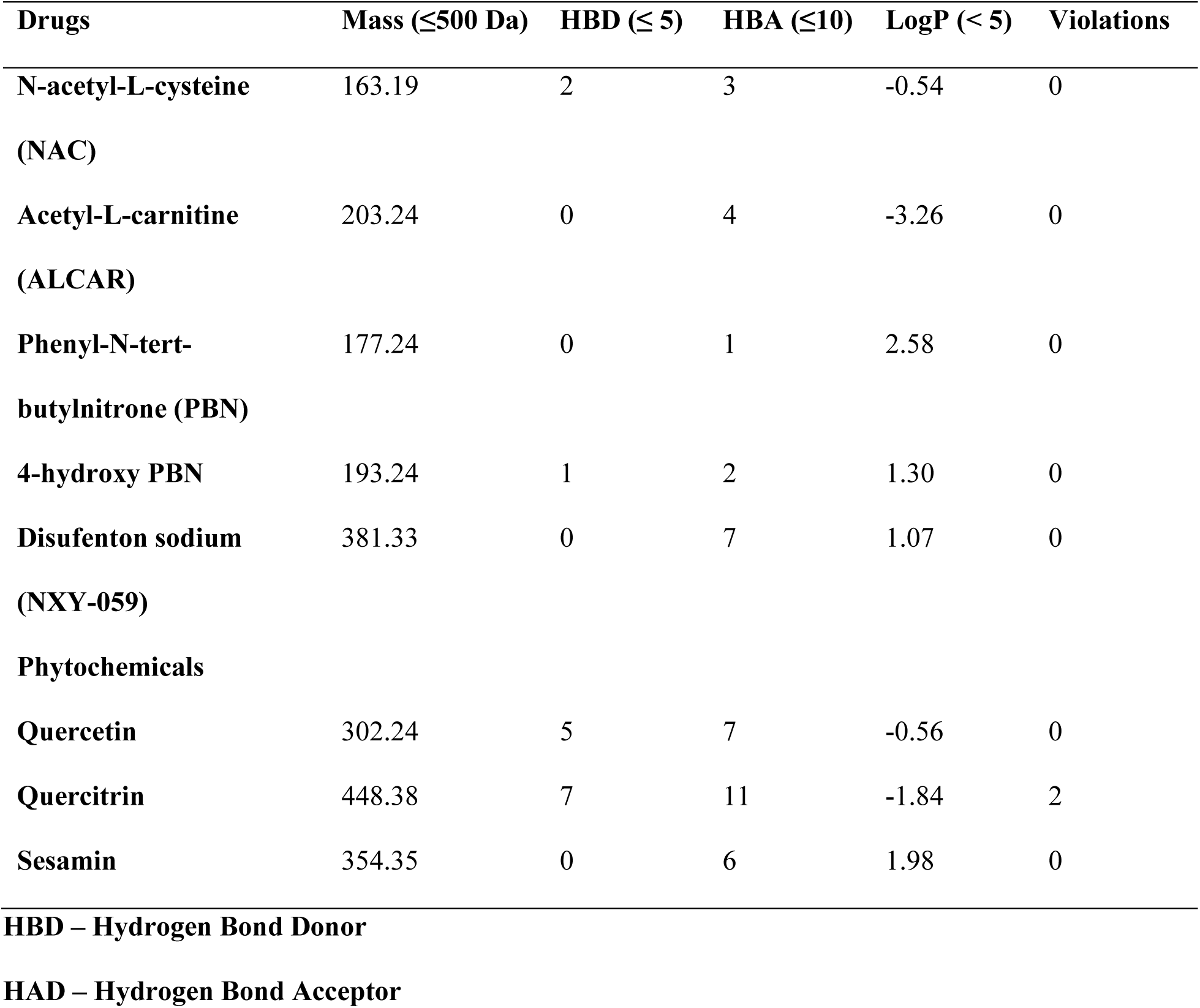
Lipinski’s Rule of Five (RO5) for to hit phytochemicals.

**Table 3:**
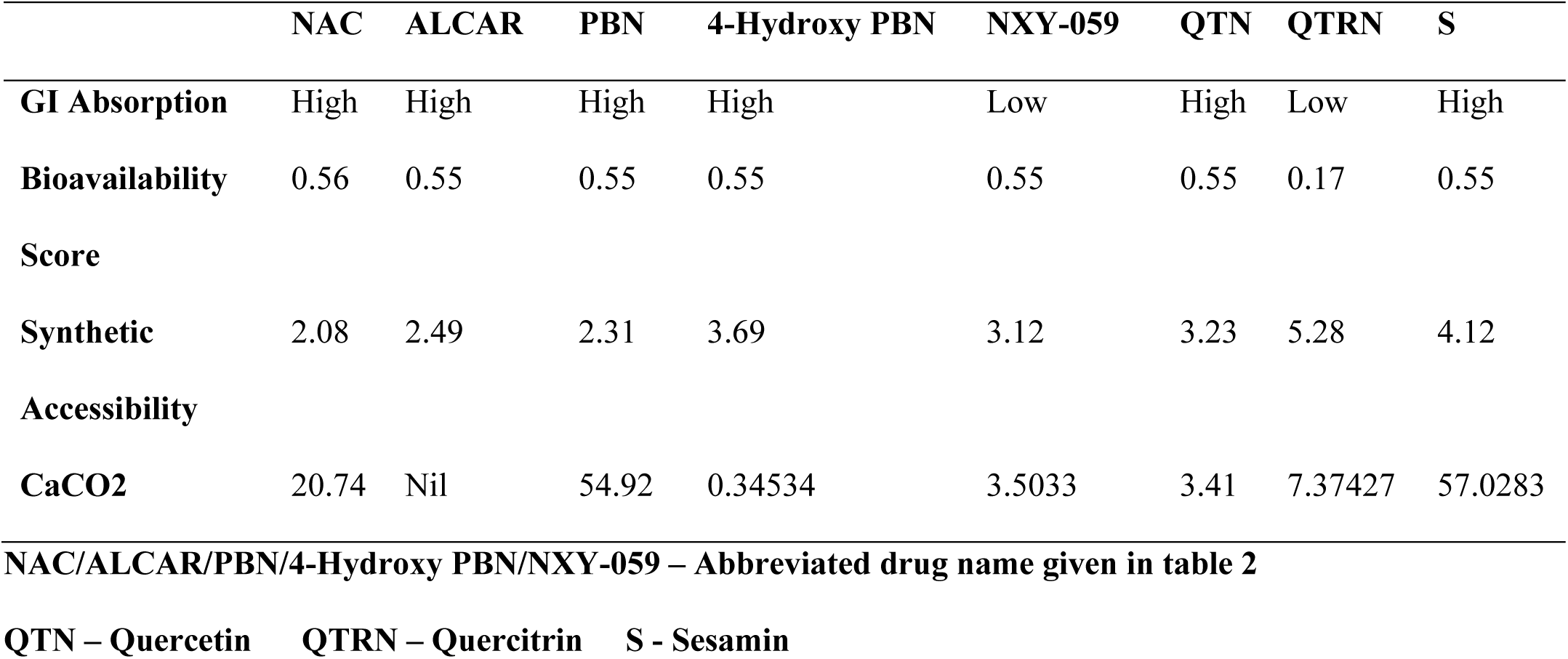
ADME and Toxicity analysis of the commercial drugs and top hit phytochemicals.

When assessing a compound’s potential to be a medication, the lipophilicity (LogP) is a crucial element to take into account. The pace at which medication molecules are absorbed by the body is influenced by LogP values. Drugs that are hydrophobic and non-polar typically have higher LogP values. In drug development, high lipophilicity is frequently preferred, especially for intracellular medicines. However, a drug’s most attractive qualities will vary depending on a number of variables. The LogP values for the top hit phytochemicals depicted a good range of values (Table 2) as compared to the commercial antioxidants, such as, Vitamin E (LogP = 6.14), Masoprocol (LogP = 2.74).

The Synthetic Accessibility (SA) rating of a substance indicates how simple it is to synthesise. A SA value of 1 indicates that the compound may be made readily, while a number of 10 indicates that the synthesis may be difficult. The phytochemicals’ SA values fell between the ranges of 3.23 and 5.28, indicating a moderate synthesis process. The quantity of the drug that will reach the desired place after injection into the body is indicated by the Bioavailability Score (BS). With a Bioavailability Score of 0.55, Quercetin and Sesamin performed well. On the other side, Quercitrin had a Bioavailability Score of 0.17, which indicated a weak reach towards the objective.

### Toxicity

The toxicity analysis was carried out to understand the initial characteristic of the top hit phytochemicals in terms of the toxicity parameters of the ligand molecules (Fig. 12). LD50 parameters were calculated for the commercial drugs and the top hit phytochemicals. The term “LD50” refers to an estimate of the amount of poison that, in the presence of control variables, will be a lethal dosage to 50% of a large number of test animals of a specific species. The value is usually expressed in milligrams of the substance being tested per kilogram of animal body weight (mg/kg). After analysing the LD50 values of the drugs and top hit phytochemicals (Table S3), it was found that Quercetin exhibited a predicted LD50 value of 150 mg/kg suggesting that a small dose of Quercetin will be toxic for the 50% population. On the other hand, Sesamin and Quercitrin showed a LD50 value of 1500 mg/kg and 5000 mg/kg respectively, which lied in the range of LD50 values exhibited by 4-Hydroxy PBN, NXY-059, NAC and PBN suggesting its potent role as an anti-oxidant along with a good MD simulation and Docking results.

**Fig. 12:**
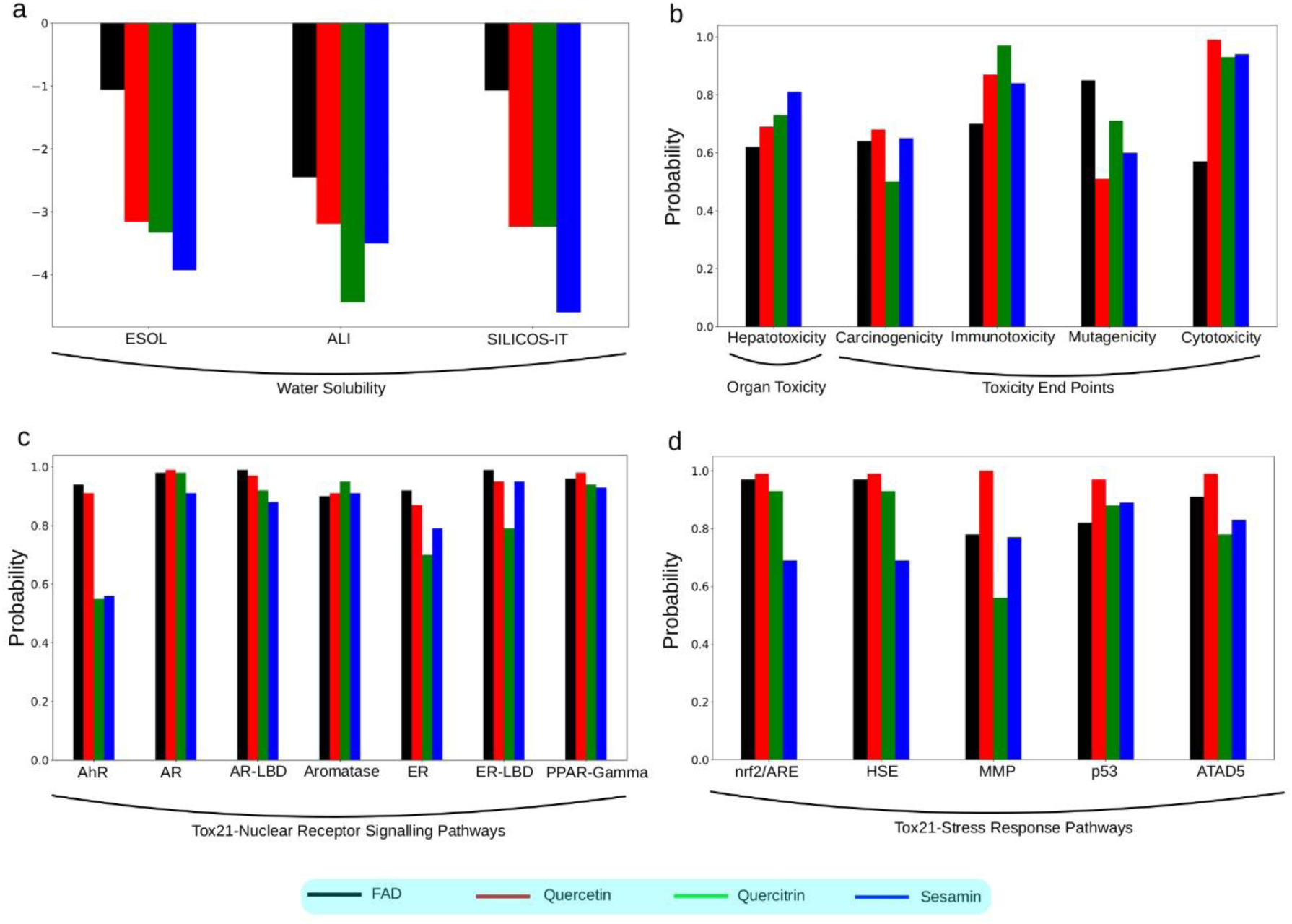
Toxicity studies for the top hit phytochemicals and FAD. (a) Water solubility of the respective ligands parameterising them for their quality solubility (b) Organ toxicity and Toxicity end points for the respective ligands quantifying them for the respective toxicity (c) Response of ligands towards Tox21-Nuclear Receptor Signalling Pathways (d) Response of respective ligands towards Tox21-Stress Response Pathways. The abbreviation are as follows: *ESOL: Estimated Solubility; AhR: Aryl Hydrocarbon Receptor; AR: Androgen Receptor; AR-LBD: Androgen Receptor Ligand Binding Domain; ER: Estrogen Receptor Alpha; ER-LBD: Estrogen Receptor Ligand Binding Domain; PPAR-Gamma: Peroxisome Proliferator Activated Receptor Gamma; nrf2/ARE: Nuclear Factor (erythroid-derived-2)-like 2/antioxidant responsive element; HSE: Heat Shock Factor Response Element; MMP: Mitochondrial Membrane Potential; p53:*

### Phosphoprotein (Tumour Suppressor); ATAD5: ATPase family AAA Domain-containing protein 5

Protox-II server was used to calculate the other relevant toxicity of the ligands. It was found that the top hit phytochemicals showed a value in the range of commercial drugs but were tagged as carcinogenic and immunogenic active by the server (Table S4). The server classified all the top hit phytochemicals (except Quercetin) as mutagenic inactive and all the top hit phytochemicals as Cytotoxic inactive. Moreover, on analysing the response of the top hit phytochemicals on the nuclear receptor signalling pathways, it was found that the server classified Sesamin as inactive in all the categories (Table S5). Quercitrin was classified as inactive in all the categories except for Aryl Hydrocarbon Receptor toxic. Furthermore, the server classified Sesamin and Quercitrin both as inactive in all the categories of Stress Response Pathways. Whereas, Quercetin was classified as toxic for Mitochondrial Membrane Potential (Table S6).

## Conclusion

The enzymatic system comprising Glutathione S-transferase (GST) and Glutathione Peroxidase (GPx) plays a pivotal role in utilizing the cellular reduced glutathione (GSH) reservoir for catalysing detoxification and antioxidation processes. In physiological contexts, GSH primarily functions to counteract cancer by neutralizing xenobiotics and oxidative agents. Augmenting glutathione levels, along with associated enzyme activity, may inadvertently diminish treatment effectiveness by scavenging these radical species. Conversely, under pathological conditions, afflicted cells adeptly exploit the glutathione system to ensure survival. Hence, combatting diseases such as malaria and cancer necessitates targeting the Glutathione Reductase (GR) enzyme to curtail GSH production. Empirical evidence underscores the feasibility of enhancing cancer therapy through precise modulation of the glutathione system. Notably, certain synthetic and natural agents exhibit the capacity to disrupt glutathione dynamics and enzyme function. This study underscores the imperative of inhibiting GR as a therapeutic cornerstone, underlining its potential impact in battling diverse diseases, and highlights the multifaceted potential of our investigation. (Narayankutty et al., 2019).

The primary objective of this study is to comprehend the antioxidative mechanism exhibited by phytochemicals derived from *Houttuynia cordata*. These compounds, potent in modulating the activity of GR enzymes across varied pathological conditions, were investigated. Our investigation identified three prominent phytochemicals—Quercetin, Quercitrin, and Sesamin— as robust interactors with the active site binding pocket of the GR protein. These compounds displayed elevated binding affinities, particularly engaging critical residues including Lys66, Thr339, Leu337, Ser177, Thr57, Cys63, Leu338, and Ile198 (Table 1), accompanied by hydrophobic interactions. Quantitative metrics such as binding scores and energies highlighted their superior binding performances compared to other phytochemicals, positioning them as prime candidates for unravelling dynamic interaction nuances. Employing a rigorous 150 ns Molecular Dynamics simulation, we discerned enhanced binding dynamics for all three compounds with GR. Notably, comparative Root Mean Square Deviation (RMSD) profiles were observed (Fig. 7). Further investigation encompassing Root Mean Square Fluctuation (RMSF) (Fig. 8a), Radius of Gyration (Fig. 9), Hydrogen Bond dynamics (Fig. 8b), and Solvent Accessible Surface Area (SASA) (Fig. 10) corroborated the slightly superior performance of Quercitrin and Sesamin in contrast to Quercetin. However, it is pivotal to emphasize that these findings, while suggestive of Quercitrin and Sesamin’s potential, do not discount Quercetin’s significance as an antioxidant candidate. Further experimental validation is imperative to ascertain their actual activity. This study unveils a comprehensive understanding of the intricate interaction dynamics, paving the way for subsequent investigations and clinical explorations. Furthermore, an in-depth Dynamic Cross-Correlation Matrix (DCCM) analysis elucidated the intricate residual dynamics of GR upon binding with the respective ligands within the active pocket. Remarkably, specific domains within GR exhibited discernible patterns of positive and negative correlation, exemplified by residues 400-420 (Fig. 11), which could potentially contribute to heightened protein stability.

In parallel, our investigation delved into the ADME-Tox domain, shedding light on the prospective therapeutic utility of these phytochemicals. Our findings underscore the intrinsic potential of *Houttuynia cordata* as a robust pharmacological resource for targeting GR enzyme activity. This revelation has profound implications, suggesting the viability of employing the plant as a natural reservoir for GR inhibition, offering a compelling alternative to conventional synthetic pharmaceutical interventions. The outcomes of this study lay the foundation for informed decisions within the pharmacological sphere, enriching the potential arsenal against pathologies reliant on GR dysregulation.

